# Temporal learning modulates post-interval ERPs in a categorization task with hidden reference durations

**DOI:** 10.1101/2022.01.25.477745

**Authors:** Mateus Silvestrin, Peter M. E. Claessens, André M. Cravo

## Abstract

The investigation of time-related activity in human electrophysiological activity has recently expanded from signals during the estimated interval to post-interval activity. Previous findings show timing-associated event-related potentials (ERPs) in both early (∼150ms) and late positive components (LPC; ∼300ms) post-interval signals. However, it is still unclear whether and what aspects of temporal information these different patterns of EEG activity are capturing, especially, if these signals are associated with interval duration or other task parameters. In the present work, we designed a modified temporal categorization task to investigate if participants’ adaptation to a changing decisional context led to changes in post-interval ERPs. Participants had to learn a hidden reference duration on each block to categorize the presented target intervals correctly. We found an early posterior N200 potential correlated to target durations and decisional context. A central LPC, which interacted with duration and learning of the hidden reference, predicted behavioral responses. Our findings add to the current evidence that interval duration modulates post-interval activity in timing categorization tasks and shows that late signals are associated with interval duration and decisional context.

## 1 Introduction

Several electrophysiological correlates have been proposed as indexes of interval duration in humans (for a review see Kononowicz et al., 2018). For many years, research focused primarily on neural signatures during the to-be-timed interval, specifically on the contingent negative variation (CNV; Van Rijn et al., 2011). Recently, an increasing number of studies have started to evaluate correlations locked to the end of target interval (Bueno and Cravo, 2021; Kruijne et al., 2021; Kononowicz and Rijn, 2014). These studies have found that early electroencephalographic (EEG) activity evoked both in sensory-related sensors and later decisional-related activity are modulated by the interval duration to be judged (Bueno and Cravo, 2021; Kruijne et al., 2021; Kononowicz and Rijn, 2014).

The majority of these studies used a temporal categorization task, in which participants judge whether a target interval is longer or shorter than a given reference interval. For instance, Kononowicz and Rijn (2014) used a categorization task where the participants were informed that there were six possible target durations (three shorter and three longer than the reference). The authors found an auditory N1P2 effect with an amplitude that increased as a function of the temporal distance from the reference interval, leading to a V-shaped pattern between the duration of the interval and EEG amplitude.

Recently, two studies have also found evidence of an early post-interval modulation of activity associated with temporal processing. Kruijne et al. (2021) used an interval categorization task where reference duration was shown (and changed) every trial and target duration was presented after a memory delay. The authors found that the P200 for the target duration showed higher amplitudes the longer the interval duration and partially predicted behavioral responses. Bueno and Cravo (2021) used multivariate pattern analyses to disentangle signal modulations associated with interval duration and task type. In their study, the authors observed an early signal modulation associated with target interval duration, similar to a P200. However, its scalp topography was more posterior, its presence was independent of time being relevant to the task, and it showed no consistent association to behavioral responses Bueno and Cravo (2021).

The pattern of findings for late event related potentials (ERPs), such as the P300 or the *Late Positive Component* (LPC), is also unclear. For instance, in their study, Kononowicz and Rijn (2014) did not find late decision-related potentials associated with temporal processing. Contrasting again the recent works of Kruijne et al. (2021) and Bueno and Cravo (2021), both found signals modulated by interval duration. On both accounts, shorter durations were associated with higher signal amplitude. However, the late signal predicted behavioral responses only in the second report.

Although temporal categorization tasks have brought significant advances in our understanding of temporal processing, this task has some shortcomings. Most studies use a single temporal reference, making it difficult to dissociate whether brain activity evoked by a given duration is associated with the actual duration of the target or with its category (longer/shorter). Moreover, participants can learn the distribution of the intervals used in the task, making it hard to dissociate whether different ERP correlates are associated with task-specific information or with more general temporal information. One possibility to overcome some of these difficulties is to investigate how neural correlates of temporal processing change as a function of participants learning and updating a specific temporal reference. Observing these changes can help understand how different aspects of temporal processing modulate behavior and brain activity.

In the present study, we developed a task where the same set of target intervals was presented in every block while the decisional context (the reference interval) changed. Importantly, participants had to learn the context of every block in the form of a hidden reference. This design allowed us to observe participants learning a new temporal reference and investigate how this learning reflected in post-interval EEG activity.

## 2 Results

### 2.1 Behavior

Twenty volunteers performed a modified interval categorization task. In each trial, a white circle (diameter 3° VA) was presented at the center of the screen for a duration equal to one of six target intervals (0.5, 0.8, 1.1, 1.4, 1.7, 2s), and participants had to judge if it was longer or shorter than the reference for that block. Contrary to regular categorization tasks, the reference interval was never presented, and participants had to learn it from feedback received at each trial. Feedback was presented immediately after the response. There were three possible reference durations: 0.65 (Short), 1.1 (Medium), and 1.45s (Long). Each block consisted of 24 trials (each interval was presented four times in total) for a total of 36 blocks (12 for each hidden reference). Critically, each target duration was shown twice in each block half to facilitate comparisons within blocks.

Behavioral performance was measured by fitting psychometric curves on each condition and block half and comparing points of subjective equality (PSE). In the current context, we stipulate the ‘point of subjective equality’ to be the interval which, according to the fit psychometric function, leads to 50% ‘longer’ responses, taking into account guess and lapse rate. We take this parameter to be the observer’s estimate of the hidden reference. As shown in (figure 2A), there are strong differences of response patterns for each reference already at the beginning of each block, and this difference increased throughout each block. A Repeated-Measures ANOVA on the estimated PSEs (figure 2B) with factors Reference (three levels: Short/Medium/Long) and Block Half (two levels: First Half and Second Half) showed a main effect of Reference (*F*[1.33, 25.08] = 253; p < 0.001; *ω*^2^ = 0.90), a main effect of Block Half (*F*[1, 19] = 4.94, p = 0.039; *ω*^2^ = 0.08) and an interaction (*F*[2, 34.22] = 63.91, p < 0.001; *ω*^2^ = 0.54). A simple main effects analysis of Block Half showed that changes in PSE are present for short (*F*[1] = 11.19, p < 0.01) and long (*F*[1] = 98.31, p < 0.001), but not for the medium reference (*F*[1] = 1.16, p = 0.295). This suggests that participants started the blocks assuming the Medium reference to be correct, and adjusted their responses according to feedbacks.

We further analyzed performance by estimating the just-noticeable difference (JND) for each reference and block. A similar ANOVA exhibited a main effect of Reference (*F*[1.61, 30.67] = 11.58; p <0.001; *ω*^2^ = 0.13), no main effect for Block Half (*F*[1, 19] = 0.56; p = 0.46) and an interaction effect (*F*[2, 34.71] = 4.45, p = 0.02, *ω*^2^ = 0.02). A simple main effects analysis of Block Half showed only the effect for the Short reference approaching significance (*F*[1] = 4.04, p = 0.059; Medium: *F*[1] = 0.054, p = 0.47;Long: *F*[1] = 2.51, p = 0.13). However, while for Short reference durations the JND decreased as the block progressed, for the Long duration, JNDs increased, in agreement with the scalar property of time.

To probe the duration of the reference participants used in each block, they were asked to generate an interval with the same duration as the hidden reference. At the end of each block, a blue circle appeared in the center of the screen, and participants had to interrupt its presentation when the duration of the reference they were using elapsed (three trials per block). Interval durations produced by participants (figure 2D) increased as reference duration increased. A repeated-measures ANOVA with factor Reference (three levels) was significant (*F*[1.23] = 78.48; p < 0.001; *ω*^2^ = 0.33). Post-hoc comparisons with Holm corrections showed differences among all comparisons (ps < 0.001), suggesting the productions were in line with the hidden reference durations.

Taken together, psychometric and production results suggest that participants discriminated the hidden references well and adjusted their performance accordingly.

### 2.2 EEG

In a first step, timing-related ERPs were identified on epochs locked to the offset of the interval (see Methods for details). We found two spatiotemporal patterns of EEG activity modulated by target duration/category in offset-locked epochs (figure 3). A first pattern with an early posterior distribution (150-250ms; P7/P8, P5/P6, PO7/PO8) which resembles an N200; and a second pattern with late fronto-central distribution (300-400ms; F1/Fz/F2, FC1/FC2, C2/Cz/C1), similar to previously reported late positive components (LPC).

To investigate whether and how hidden references modulated these two ERPs, we applied a repeated-measures ANOVA with factors Target Duration (six levels) and Reference (three levels). The early ERP showed both main effects of Target Duration (*F*[2.13, 40.50] = 50.95, p < 0.001, *ω*^2^ = 0.39) and Reference (*F*[1.53, 29.15] = 4.21, p = 0.03, *ω*^2^ = 0.007) and no interaction (*F*[6.36, 120.87] = 1.18, p = 0.32). The reference effect showed a linear trend across short, medium and long reference (*β* = −0.21, t = -2.77, p = 0.009). The Target Duration effects were inspected with a Helmert contrast. The ERP amplitudes for each of the first three durations (500, 800 and 1100ms) differed from the aggregated means of the subsequent ones (all corrected p-values < 0.04).

For the LPC, the overall pattern of results figure 3D showed higher amplitudes for shorter target durations and subtle differences in amplitudes between references in intermediate durations. We found a main effect of Target Duration (*F*[1.76, 34] = 64.26; p < 0.001, *ω*^2^ = 0.58), a main effect for Reference (*F*[2, 38] = 3.64; p = 0.04, *ω*^2^ = 0.01) and an interaction (*F*[10, 190] = 6.46, p = 0.004, *ω*^2^ = 0.02). We investigated the interaction with a simple main-effects analysis for reference, which showed differences in amplitude among references for target durations of 800ms (*F*[2] = 4.44; p = 0.02) and 1100ms (*F*[2] = 9.67, p <0.001, all other p > 0.26).

### 2.3 Evolution of post-interval EEG activity as function of learning

To explore further how the effect of reference evolved during each block, we separated each block into two halves. Our task design defined that each target duration appeared twice each block half to investigate this learning effect.

For the early ERP, data from all intervals were included, given that in the first analysis, we found only the main results of duration and reference. In an ANOVA with factors Target Duration (six levels), Reference (three levels) and Block Half (two levels), we found a main effect for Target Duration (*F*[2.13, 40.45] = 51, p <0.001, *ω*^2^ = 0.39) and a main effect for Reference (*F*[1.53, 29.11] = 4.1, p = 0.04, *ω*^2^ = 0.01). The effect of Block Half, however, was not significant (*F*[1, 19] = 0.02, p = 0.9). None of the interactions were significant (ps > 0.29, see OSF repository for complete ANOVA table). Target Duration effects were investigated with a Helmert contrast: 0.5, 0.8, and 1.1s target durations were significantly lower than their subsequent levels (ps <0.04; see the OSF repository for the complete table). Reference effects showed a significant negative linear trend (*β* = −0.21, t = -2.72, p = 0.01).

For the LPC, we focused on target durations of 0.8s and 1.1s, given that only these two were significantly modulated by the hidden block reference. The average amplitudes of these ERPs were submitted to a repeated-measures ANOVA with factors Reference (three levels: short/medium/long), Target Duration (two levels: 800 and 1100ms), and Block Half (two levels). Consistent with our previous findings, there was a main effect of Target Duration (*F*[1, 19] = 37.74, p <0.001, *ω*^2^ = 0.17), Reference (*F*[2, 38] = 63.24, p <0.001, *ω*^2^ = 0.06) and a main effect of Block Half (*F*[1, 19] = 4.66, p = 0.04, *ω*^2^ = 0.02). Our main interest was in the interaction of block half and reference, which could reveal if the effect of the hidden reference emerged only on the second half. We found a small effect for this interaction (*F*[2, 38] = 3.31, p = 0.047, *ω*^2^ = 0.01). A simple-main effects analysis for the effect of reference on each block half showed that the effect of reference was not significant on the first half, (*F*[2] = 1.44, p = 0.25), but was on the second half (*F*[2] = 13.55, p <0.001). All other effects were not significant (ps > 0.06, see OSF repository for complete ANOVA table).

The results presented so far suggest that ERP amplitude indexed mixed information about both target duration and target category relative to the hidden reference. Therefore, in the last analysis, we investigated whether ERP amplitude predicted participants’ responses on a trial-by-trial basis. We focused on the two reported ERPs (the parietal N200 and the LPC). We ran a generalized linear model per participant with the predictors Target Duration and Normalized ERP Amplitude (the EEG variability not accounted by either target duration or by reference; see Methods). Our results indicated that, for the early N200, only Target Duration was correlated with response (target duration: t(19) = 19.85, p<0.001, Cohen’s D = 4.44; ERP: t(19) = 1.33, p = 0.20). For the LPC, both variables were correlated with participants’ responses (target duration: t(19) = 19.48, p <0.001, Cohen’s D = 4.30; ERP: t(19) = -4.24, p <0.001, Cohen’s D = 0.95).

## 3 Discussion

In this study, we investigated the neural signatures of a time categorization task with hidden references. The task was designed to examine the neural correlates before and after learning a hidden reference. Our behavioral results showed that hidden temporal references can be learned effectively in just a few trials, as shown by the psychophysical parameters of the categorization and production tasks. Our EEG findings showed that activity at the offset was correlated with temporal processing. The two main patterns of EEG activity modulated by time were an early N200 and an LPC.

An early posterior N200 was present and modulated by time and reference in our findings. The modulation of this ERP by duration was present up to 1.1s, after which its amplitude seemed to plateau (figure 3C). This N200 was also modulated by reference, with the longer reference shifting down its amplitude. This pattern could reflect the N200 indexing how much shorter the current interval is relative to the reference, explaining its stabilization after intermediate durations. On the other hand, the effect seems to be driven strongly by the shortest interval and did not, in our findings, correlate with behavior. Lastly, the N200 was also not modulated by how well participants knew the reference, with similar results during the two halves of the blocks.

One of the first studies that looked carefully at post-interval activity showed an N1P2 that was modulated by target durations (Kononowicz and Rijn, 2014). In their study, Kononowicz and Rijn (2014) found a V-shaped N1P2, with higher amplitudes for intervals further from the reference, irrespective of whether the target was shorter or longer than the reference. Recently, Kruijne et al. (2021) showed a fronto-central P200 modulated by duration and subjective perception, with higher amplitudes in longer durations and higher chances to answer accordingly. Finally, in a recent study, Bueno and Cravo (2021) found an early posterior activity modulated by target duration, with a time course also similar to a P200, but with a more parieto-occipital topography than Kruijne et al. (2021). As can be seen, although the current experiment and all three studies (Kononowicz and Rijn, 2014; Kruijne et al., 2021; Bueno and Cravo, 2021) have found an early modulation of EEG activity by time, each found a different pattern of results. Due to the marked differences in task designs, connecting these effects in early components is challenging, and differences might have emerged by the stimuli used (auditory or visual), by whether the interval was filled (marked by the onset-offset of the target) or empty (marked by two onsets). However, it does seem that, although several studies have found early modulation of EEG activity, which activity is modulated and how it is modulated depends heavily on task particularities and demands.

For the LPC activity, we found a strong modulation of interval durations, similar to previous results (Bueno and Cravo, 2021; Kruijne et al., 2021). This EEG activity had a higher amplitude for shorter durations and stabilized after intermediate durations. Unlike the N200, the hidden reference modulation was restricted to specific intervals. Intervals that did not change category (such as 0.5s, which was always shorter than the reference, or 2s, which was always longer) were not modulated when participants learned the hidden reference. This result is important because it shows that the LPC does not seem to be indexing only how much shorter a given interval is from the reference. At the same time, it does show that the LPC is being modulated by the temporal reference and cannot be explained by a surprise due to interval ending.

This pattern of LPC amplitudes has been found previously in categorization task experiments with explicit reference intervals (Bannier et al., 2019; Bueno and Cravo, 2021; Kruijne et al., 2021; Wiener and Thompson, 2015) ^1^. In these studies, the general finding was of amplitudes being larger as a function of how much shorter the target interval is relative to the reference, up to reference interval, after which the LPC plateaus (Bannier et al., 2019; Bueno and Cravo, 2021; Kruijne et al., 2021). Previous studies have shown that this pattern is present only when duration is task-relevant (Bueno and Cravo, 2021) and for both bisection and generalization tasks (Bannier et al., 2019), with the effect of duration being stronger in bisection tasks (Bannier et al., 2019). However, the term LPC in the timing literature has been used for a wide range of activity found in different sensors and time-periods (Wiener and Thompson, 2015; Gontier et al., 2007, 2009; Paul et al., 2011). Moreover, it is unclear whether and how the LPC differs from other late-decisional components, such as the P300 (Polich, 2007). Thus, it is hard to reconcile all previous studies that have claimed to have found a correlation between the LPC and temporal processing. On the other hand, our current findings are consistent with previous results showing a late frontal-central component that is correlated with temporal processing (Bannier et al., 2019; Bueno and Cravo, 2021; Kruijne et al., 2021).

A two-stage drift-diffusion model has been proposed for time categorization tasks (Simen et al., 2011; Balci and Simen, 2014, 2016). The first stage is dedicated to timing accumulation during the interval. In contrast, in the second stage, which starts at the interval offset, evidence is accumulated towards one of two decision thresholds (“short” and “long” responses). Given that our EEG results were mainly at the interval offset, it is natural to examine whether the LPC signal might be associated with some of the decision process parameters. One possibility is that the LPC amplitude might serve as a signature of the drift rate towards the “short response” threshold at the decision phase. In their model, Balci and Simen (2014) propose that the drift rate at the second stage is correlated with the difference between the starting point and the midpoint between thresholds. In this view, the drift rate is maximum at the shortest and longest duration and should be lower in intermediate durations. This is partially consistent with the pattern of the LPC amplitude, which is correlated only monotonically with interval duration. Critically, in the model, the effect of different references leads to different drift rates for the same physical durations in the second stage due to where the first stage ends. Specifically, the prediction is to obtain higher drift rates for the same duration when the reference is longer, which we observe in the LPC: the same physical duration has a higher LPC when the reference is longer. However, in our results, LPC amplitudes are modulated by the reference only for intermediate durations that change category (shorter/longer) for different references. Whether this pattern is compatible with the proposed model is unclear, and can depend on how the thresholds for the short/long responses are fixed.

Contrary to the two-stage model, we do not see substantial amplitude differences in the LPC for longer durations. This might suggest that the first stage uses a single threshold as the reference interval, and if the accumulation in the first stage crosses this threshold, no second stage is needed. Although this proposal is contrary to the original model, in which a decision should always occur during the second stage (Simen et al., 2011; Balci and Simen, 2014, 2016), it is consistent with other LPC results in temporal tasks (Bannier et al., 2019; Bueno and Cravo, 2021; Kruijne et al., 2021) and with previous behavioral and modeling results (Wiener et al., 2018).

The idea of the LPC reflecting the drift rate of the second stage of the decisional model is consistent with recent studies in decision making. It has been proposed that processes of the drift diffusion models are reflected in late ERP, such as the P300 (Twomey et al., 2015). However, it is usually assumed that the drift rate should reflect on the ramp-up of the P300 and not on its amplitude, making it currently unclear why the differences found in temporal decisions, so far, are reflected on the amplitude of the LPC.

An important limitation in our design was that only some of the presented intervals changed their category of shorter/longer as a function of learning. In our experiment, each interval duration was shown four times per block, giving us a reasonable number of trials to detect electrophysiological changes in the dynamic decisional context. However, it also restricted the number of intervals that changed its category of shorter/longer as a function of the hidden reference. Future studies with dynamic categorization contexts can maximize the number of intervals within different categories across different contexts.

In conclusion, our findings add to the current evidence that post-interval activity is modulated by interval duration in timing tasks. This activity changed as a function of the temporal reference and its learning. These findings suggest that late EEG activity can be used to track how timing is processed and, combined with future tasks, help compare predictions from different theoretical models of temporal processing.

## 4 Methods

### 4.1 Participants

Twenty healthy adults (24*±*4 years), undergraduate and graduate students with reported normal or corrected to normal vision participated in the study. All participants filled informed consent forms before the experiment. UFABC’s Ethical Committee approved the research protocol.

### 4.2 Task design

An interval categorization task was modified to include dynamical learning of hidden reference intervals. The task was designed in Matlab (MATLAB, 2014) with Psychtoolbox (Brainard, 1997) and presented on a CRT screen (refresh rate = 60Hz). Participants had to categorize target durations as shorter or longer than a hidden reference interval in each trial. Feedback was given immediately after the response. Importantly, the hidden reference was never presented to participants; they had to learn it based on feedbacks as the block progressed. Participants were not informed how many possible references existed. By the end of the block, participants were asked to report the hidden interval duration by producing it three times. The percentage of correct responses in the categorization task was also shown at the end of the block as overall feedback. Figure 1 shows the experiment design.

**Figure 1:**
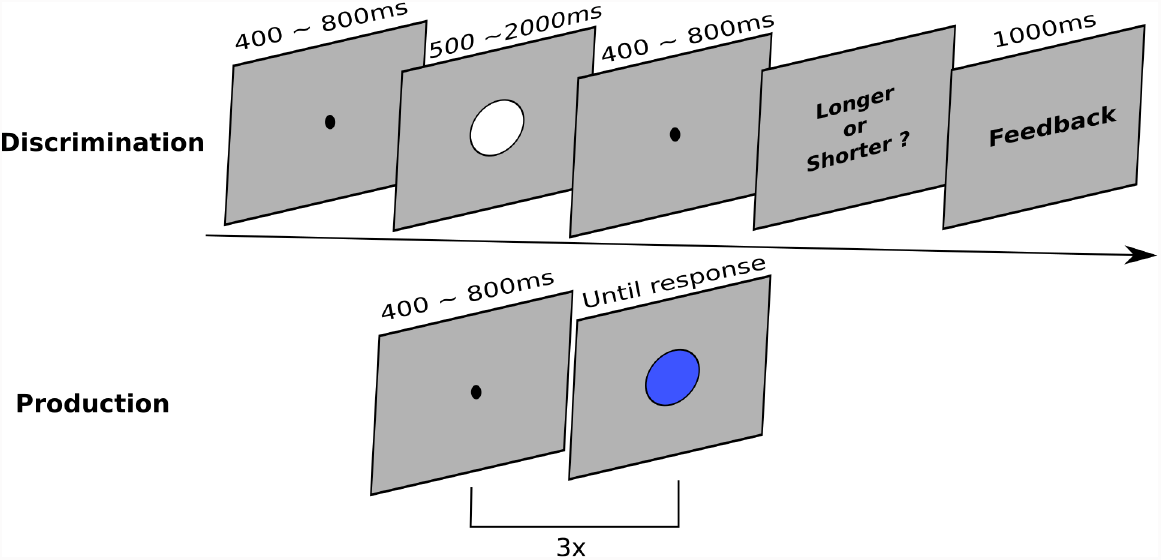
Experimental design. Upper panel: In each trial, a target duration was presented as a circle in the center of the screen (0.5, 0.8, 1.1, 1.4, 1.7, or 2s). Participants had to judge whether it was longer or shorter than the hidden reference for that block (Short:0.65, Medium:1.1, Long:1.45s). Lower panel: at the end of each block, participants were asked to produce intervals with the same duration as the hidden reference.

**Figure 2:**
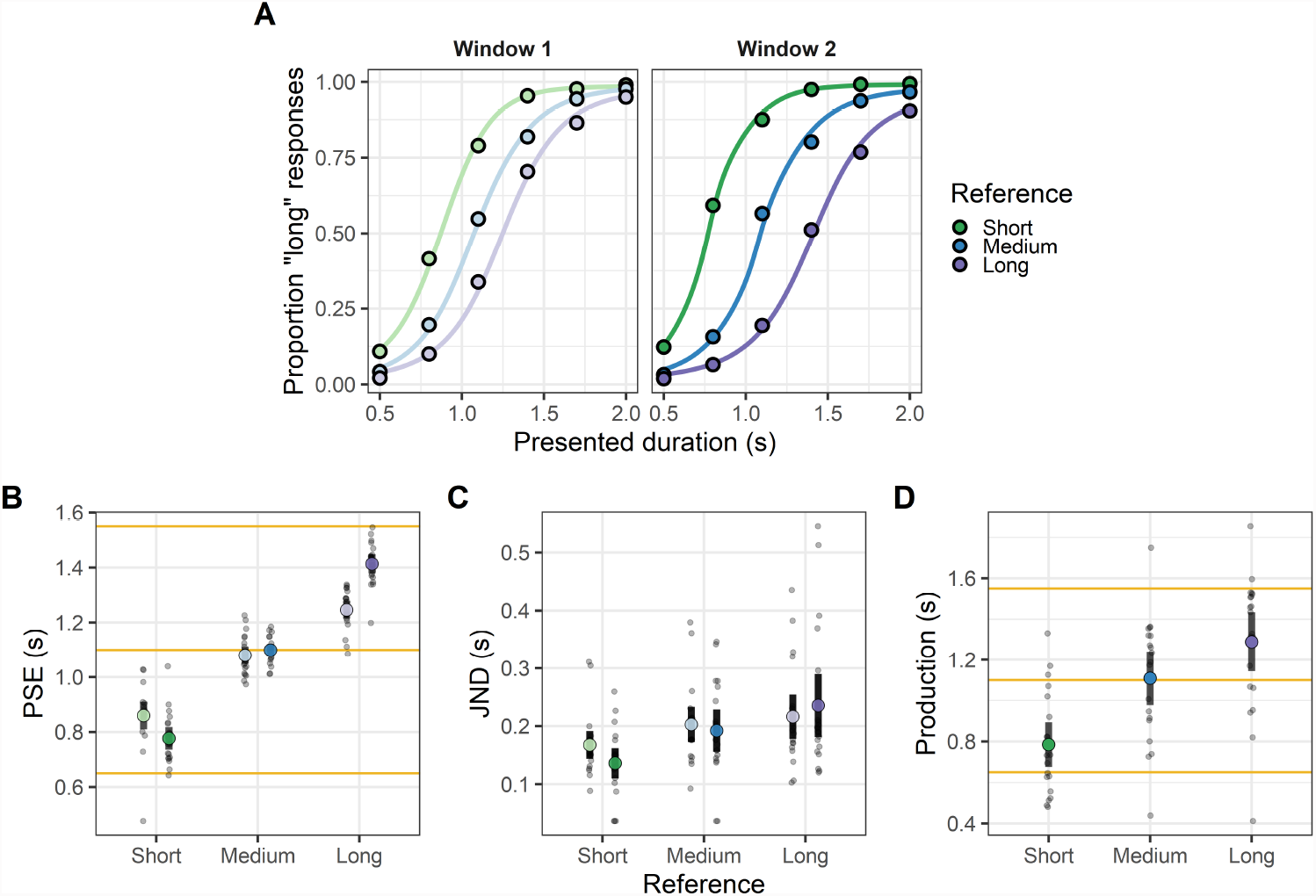
Behavioral results. (A) Average psychometric curves for the first (left panel) and the second (right panel) half within each block. Grey dots show individual averages. (B) Points of subjective equality-PSE (mean ± s.e.m.) across tasks and halves (light gray dots show individual data, yellow horizontal lines show reference positions). (C) Just noticeable difference-JND across references and halves. (D) Average interval productions at the end of the blocks.

**Figure 3:**
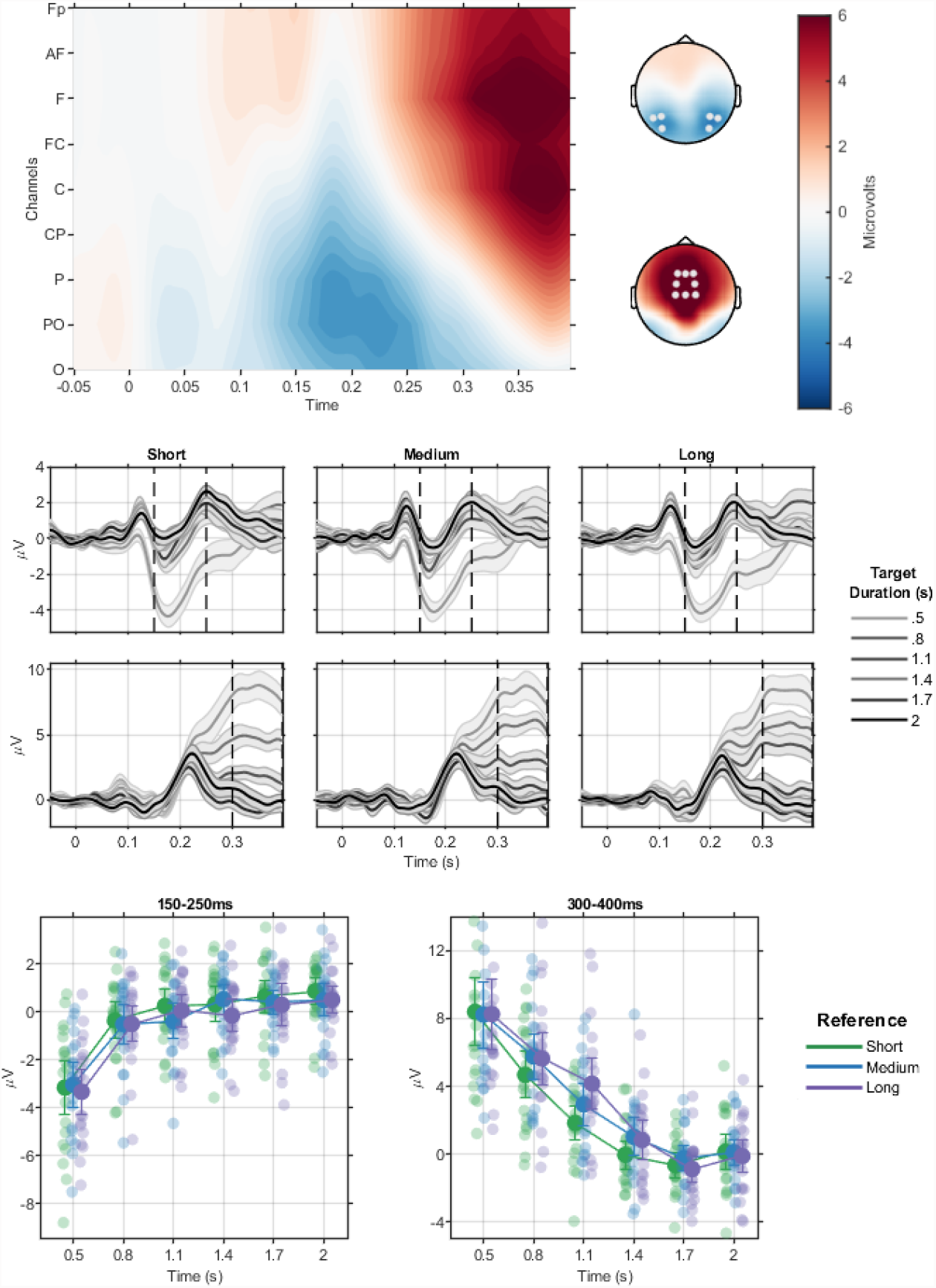
Target-offset-locked ERPs. Zero in time axes represent target offset. (A) Signal amplitudes for the difference between trials in the shortest (0.5s) and longest (2s) target durations for all electrodes and time points. The Y axis shows electrodes in a posterior-anterior sequence; the names of most electrodes are omitted for ease of visualization. The topographies show the scalp distributions for the early (150-250ms, top) and late (300-400ms, bottom) ERPs. Points represent chosen electrodes for each ERP. (B) ERP time-courses for target and reference, again for the early and late ERPs. Bands represent standard-error of the mean. (C) Mean ERP amplitudes for target and reference, error bars show the 95% confidence interval.

**Figure 4:**
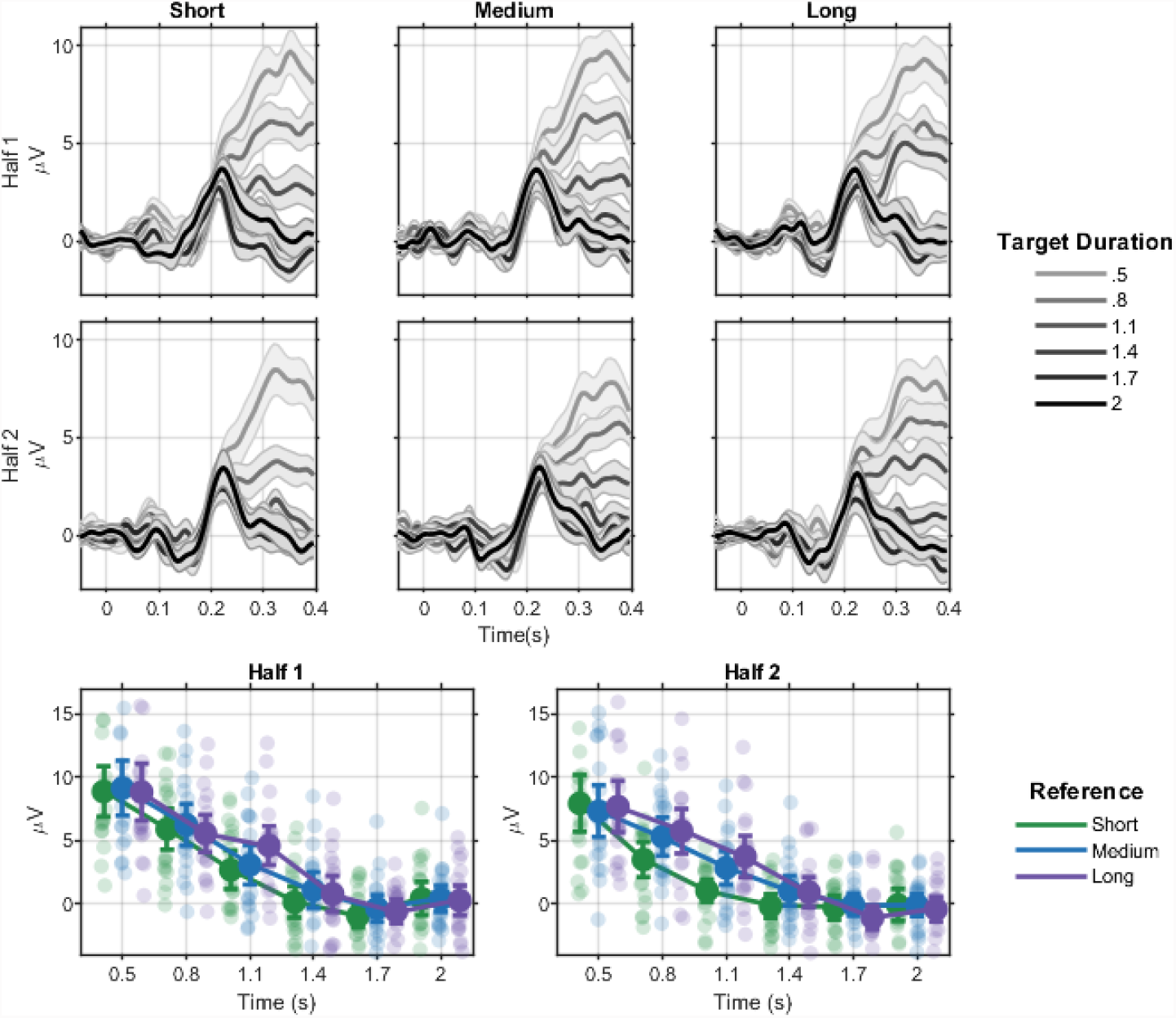
Posterior target offset ERP by block half. (A) Time-courses by target duration, reference and block half. (B) Error-bars for the period 300-400ms. Error bars show the 95% confidence interval.

The experiment had a total of 36 blocks, twelve blocks for each of the hidden reference intervals (0.65s, 1.1s, and 1.45s). Blocks were composed of twenty-four categorization trials and three production trials. Each categorization trial started with a fixation point (0.45-0.65s), then a target duration was shown as a white circle (three visual degrees diameter) that remained onscreen for one of the following durations: 0.5, 0.8, 1.1, 1.4, 1.7 or 2.0s. The presentation was pseudo-random: each duration appeared twice on each block half. We expected the first twelve presentations to allow participants to learn the hidden reference and the last twelve to show better performance due to this learning. After another fixation point (0.4-0.8s), “make judgment” appeared onscreen until a response was given. Responses were delivered with a button-press in an answer box (left button for “shorter”, right button for “longer”). If responses delay was over 2s, the trial was coded as missed. Immediately after the response (or elapsed time), feedback was shown as “hit” in green, or “miss” in red on the center of the screen for 1s. For blocks of the medium reference (1.1s), when the target duration was the same as reference duration, the feedback was always “miss”.

At the end of the block, participants produced the hidden reference they had learned. In each production trial, participants interrupted the presentation of a blue circle in the center of the screen with a button-press. There were three production trials every block (ISIs: 0.4-0.8s).

### 4.3 Behavioral Analyses

Data were preprocessed in R (version 3.5.1; R Core Team, 2021), and inferential analysis was run in JASP (version 0.14.1; JASP Team, 2021). Categorization data where participants missed the response were removed (less than 1%).

Psychometric functions were fitted to categorization data with the Quickpsy package (version 0.1.5; Linares and López-Moliner, 2016). Each block was separated into two halves of 12 trials, psychometric functions were calculated for each participant, reference and half (144 points per estimation). A logistic function was fitted with parameter boundaries (min, max): threshold (0, 2.1), slope (0, 30), guess rate (0, 0.05), lapse rate (0, 0.05). A bootstrap (n = 1000) was applied to calculate the deviance of estimated models: goodness of fit was in general high (mean across all curves and participants = 0.89, range: [0.24, 1]).

To investigate differences in psychometric parameters in different decisional contexts and as each block progressed, points of subjective equality (PSEs) and just-noticible differences (JNDs) were compared with two-way repeated-measures ANOVAs with factors Reference (3 levels) and Block Half (2 levels).

Interval production data where durations were lower than 0.25s or higher than 4s were removed (less than 1%). A one-way repeated-measures ANOVA with factor Reference (three levels) was used to investigate if productions were different across hidden references.

### 4.4 EEG

EEG data were acquired with a QuickAmp amplifier (Brainvision) at 1000Hz with 64 ActiCap electrodes in the International 10-10 system, except for the TP9 and TP10 electrodes moved from the mastoids to ear lobules. Oculogram was also recorded with two pairs of bipolar electrodes. Data were preprocessed with MNE-Python (version 0.17; Gramfort et al., 2013). Preprocessing comprised re-referencing the signal to the mean of ear electrodes, ICA ocular artifact correction, and band-pass filtering. For epochs centered at the onset of target stimuli filter was set to 0.05-30Hz, while for the epochs centered at target offset, it was set to 0.1-30Hz to minimize leakage of negative trending potentials from the target presentation period. Target onset epochs ranged from -200ms to target duration. Target offset epochs went from -200 to 400ms relative to target offset. Epochs were DC detrended, baseline-corrected by subtracting from the signal either the average amplitude of the period from -200ms to 0ms (onset epochs) or the average from -50ms to 50ms (offset epochs) and finally, epochs were resampled to 250Hz. Epochs with amplitudes higher than 15*µ*V were rejected (mean 6% per participant).

ERP analyses were developed in Matlab with the Fieldtrip Toolbox (Oostenveld et al., 2010). To choose electrodes and moments of interest, we used a collapsed localizer approach (Luck and Gaspelin, 2017): we defined regions and times of interest (ROIs and TOIs, respectively) based on ROIs and TOIs from the difference between the shortest and longest duration across all conditions. Notably, these values were also always shorter or longer than the reference, irrespective of the block. The rationale for this approach is to use extreme conditions to circumscribe ROIs and TOIs and then investigate more nuanced effects (i.e. other levels of factors) in them. Therefore, in a second step, EEG activity at these ROIs and TOIs for all durations were used to investigate how they were modulated by target duration and reference. Based on this localizer, two ERPs were selected (P8, P6, P5, P7, PO8, PO7 at 150-250ms; F1, Fz, F2, FC1, FC2, C2, Cz, C1 at 300-400ms).

We calculated the mean amplitude for each target duration for offset-locked ERPs at each reference per participant. Effects of reference duration and Target Duration on amplitudes were investigated with repeated-measures 3×6 ANOVA. Our experimental design was developed to examine the effects of learning during the blocks by presenting each target duration twice on each block half. Therefore, we investigated changes in ERP amplitudes for target durations where reference effects were significant (800 and 1100ms). We applied a three-way repeated-measures ANOVA including the factor Block Half (two levels) alongside Target Duration (two levels) and Reference (3 levels). In the absence of interactions, we further investigated main effects with Helmert or polynomial contrasts, in the presence of interactions we applied simple main-effects analyses.

We investigated whether trial-by-trial fluctuations in EEG amplitude were correlated with participants’ responses. We fitted a generalized linear model (logit link) per participant with the response (shorter/longer) as the dependent variable and target duration and single-trial ERPs as predictors. To account for the strong correlation between EEG amplitude and duration/hidden reference, for each hidden reference and duration, the average ERP amplitude was subtracted from the single-trial EEG signal. The regression coefficient of this normalized ERP and response capture whether trials in which the ERP amplitude for that duration and reference was higher/lower than the average amplitude was correlated with the probability of the participant responding longer/shorter. The regression was performed per participant and, at the group level, coefficient values were tested against zero with a t-test.

### 4.5 Code Accessibility

Task and analysis code, as well as raw data will be released on publication

## Acknowledgments

The authors would like to thank Fernanda Dantas Bueno for suggestions on an early draft of the manuscript.

## Funding

MS was supported by grant #2016/24951-2, São Paulo Research Foundation (FAPESP). AMC was supported by grant #2017/25161-8, São Paulo Research Foundation (FAPESP).

## Footnotes

1 A similar pattern was a also recently reported by Nir Ofir and Ayelet Landau in a presentation that be seen at https://www.youtube.com/watch?v=GY2OklIKyiA

## Notes

### Competing Interest Statement

The authors have declared no competing interest.

